# Gustatory cortex is involved in evidence accumulation during food choice

**DOI:** 10.1101/2021.12.21.473608

**Authors:** Ali Ataei, Arash Amini, Ali Ghazizadeh

## Abstract

Food choice is one of the most fundamental and most frequent value-based decisions for all animals including humans. However, the neural circuitry involved in food-based decisions is only recently being addressed. Given the relatively fast dynamics of decision formation, EEG-informed fMRI analysis is highly beneficial for localizing this circuitry in humans. Here by using the EEG correlates of evidence accumulation in a simultaneously recorded EEG-fMRI dataset, we found a significant role for the right temporal-parietal operculum (PO) and medial insula including gustatory cortex (GC) in binary choice between food items. These activations were uncovered by using the “EEG energy” (power 2) as the BOLD regressor and were missed if conventional analysis with the EEG signal itself were to be used, in agreement with theoretical predictions for EEG and BOLD relations. No significant positive correlations were found with higher powers of EEG (powers 3 or 4) pointing to specificity and sufficiency of EEG energy as the main correlate of the BOLD response. This finding extends the role of cortical areas traditionally involved in palatability processing to value-based decision making and offers the “EEG energy” as a key regressor of BOLD response in simultaneous EEG-fMRI designs.

## Introduction

The choice of what to eat is probably one of the most common and yet most basic forms of decision making in the Animalia kingdom. This decision-making problem like any other, requires deliberation to commit intentions in favor of one choice to the exclusion of others. In decision making with uncertain sensory information (aka perceptual decision making), this process is often modelled with a drift-diffusion process supposedly representing evidence accumulation in favor of a given option (Britten et al. 1996; Mazurek et al. 2003; Gold and Shadlen 2007; Hanks et al. 2015). Neural correlates of perceptual decision-making is found in a couple of brain areas most notably the lateral intra-parietal sulcus (LIP) and the prefrontal cortex (Gold and Shadlen 2007; Pisauro et al. 2017). The accumulation of evidence during perceptual decision making is also observed in human electroencephalography (EEG) (Philiastides et al. 2014). In value-based decision making however, the evidence accumulation is done not on the momentary external evidence but on the mnemonic internal variables representing the subjective value of choice items (Bakkour et al. 2019).

Recent studies have extended the drift diffusion model of choice to value-based decision by relating an item’s subjective value to the drift term in the decision variable (DV) (Krajbich et al. 2010; Milosavljevic et al. 2010). Given the relatively rapid evolution of DV in time, neural correlates of such a process has to be searched for by methods with sufficient temporal resolution such as single unit electrophysiology or EEG. In particular, EEG studies have found neural correlates of value-based decision making across centro-parietal electrodes reflected in the raw EEG or gamma band signals (Polanía et al. 2014; Pisauro et al. 2017). However, the low spatial resolution of EEG prevents accurate localization of brain loci for value-based evidence accumulation. One work-around is to use simultaneous EEG-fMRI which combines the localization strength of fMRI with high temporal resolution of EEG (Pisauro et al. 2017). Indeed, with this technique, (Pisauro et al. 2017) found EEG signal correlates of DV in a value-based decision-making task and then used these EEG correlates as a regressor on the BOLD responses (EEG-informed fMRI analysis) to find brain regions involved. Using this method, signatures of evidence accumulation was found in the posterior-medial prefrontal cortex (pMFC). But due to the highly nonlinear mapping from electric potentials to the BOLD signal, the raw EEG may not be the best regressor for BOLD in an EEG-informed fMRI analysis.

Here, we argue that from a theoretical standpoint based on physics of EEG and fMRI, a quadratic relation between BOLD and EEG (i.e. EEG energy) may be more accurate as a first-order approximation. This conjecture is in agreement with previous reports that suggest a linear relation between the mean power of event-related EEG sources and the neural efficacy (input of vascular system for BOLD response) (Wan et al. 2006) or between EEG energy across various frequency bands and the BOLD signal (de Munck et al. 2009; Scheeringa et al. 2009; Sato et al. 2010). Notably, reanalysis of simultaneous EEG-fMRI data recorded during a value-based decision-making task with EEG energy as the BOLD regressor, revealed significant activations in cortical areas involved in palatability processing including the insula and operculum which were missed in previous analysis with raw EEG signal.

## Materials & Methods

### Data and analysis

We used the open-access simultaneous EEG-fMRI data associated with (Pisauro et al. 2017): https://openneuro.org/datasets/ds001219/versions/1.0.0. The data was partly preprocessed. We performed the remaining proceeding preprocessing steps similar to (Pisauro et al. 2017) except those described below:

### fMRI preprocessing

We used FSL to register the cleaned EPI images to the MNI space just as in (Pisauro et al. 2017) using six-parameter rigid body transformation and the nonlinear registration tool, except for subject number 20, for whom we used 12 parameter affine transformation to map his EPI to his structural image, due to a need for scaling in this case. Then we normalized the registered EPI time series to percentage of change with respect to time average of each voxel.

### Building EEG & EEG energy regressors

We used the raw signal of the *best* EEG electrode in the decision time period as described in (Pisauro et al. 2017) (which takes zero value outside the decision periods, Figure S1-b). After convolving the EEG regressor with the HRF, we subsampled this signal in intervals equal to the fMRI repetition time (here, TR=2.5s) and replaced the signal at each TR by its temporal mean within that TR. The EEG signal in (Pisauro et al. 2017) was first subsampled to 50ms resolution, then convolved with HRF and then subsampled to fMRI TR. Our downsampling method does not result in significant difference from that in (Pisauro et al. 2017), since the signal was smoothed due to convolution with the HRF. We did the same procedure but with the square of the EEG signal to build the “EEG energy” regressor. We demeaned all regressors. For the normalized regressors (Figure 5), we also divided them by their standard deviation.

### fMRI analysis

We did the analyses in AFNI 20.2.05.

The model for each primary GLM is as:

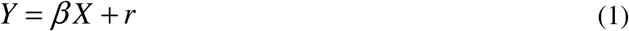

Where, *Y* is the time series of the normalized BOLD response of a single voxel for *T* time samples, X is a *n* × *T* design matrix with rows representing *n* regressors. *β* is a 1 × *n* vector, containing the regression weights for each regressor for this particular voxel and *r* is the 1 × *T* residual of this regression.

The model for each secondary GLM (second step of step-wise GLM) is as:

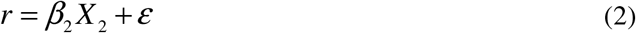

Where, r is the residual from the primary GLM for a specific voxel, *X*_2_ is the design matrix for the regressors of interest and *ϵ* is the regression residual.

We performed all our GLMs in AFNI via “3dREMLfit”. For group-level analysis, we used “3dttest++”. The group-level activation maps were then masked by the grey matter mask associated with the standard MNI brain with resolution of 2mm (Results for the raw EEG regressor were not masked to make them comparable with (Pisauro et al. 2017)). By applying 3dFWHMx on these group-level residuals, we estimated the parameters for the non-Gaussian spatial autocorrelation function of the fMRI noise. Then, using 3dClustSim, we calculated the cluster thresholds for various *p*-values such that the probability of a false positive cluster among the *p*-thresholded clusters is less than α=0.05. The inflated surfaces are presented using SUMA 20.2.05.

## Results

We re-examined the simultaneous EEG-fMRI data from (Pisauro et al. 2017), in which subjects were asked to choose between pairs of previously rated snack items and to indicate their choice with a button press (Figure 1a). The difficulty of the decision was controlled by the value difference (VD) in the ratings of the presented items. The EEG electrode that best matched “theoretical prediction of a dynamical sequential sampling model (SSM) fitted to the behavioral data of each subject” was used as a regressor against the BOLD signal in all voxels for localization of brain regions supporting decision making in this task (Figure 1b-c, see (Pisauro et al. 2017) for further details on the task and previous findings).

**Fig. 1.**
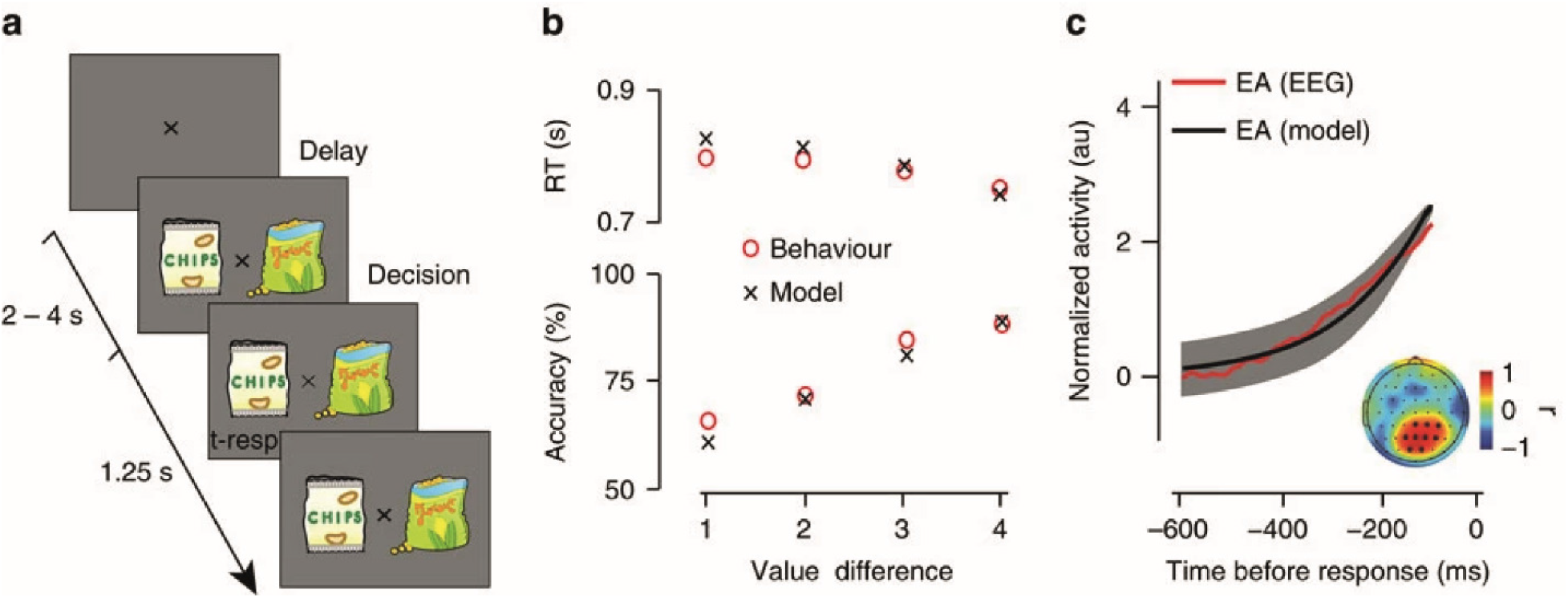
Task design, behavioral and modelling results and EEG. (a) Schematic representation of the experimental paradigm. After a variable delay (2–4 s), two stimuli (snack items) were presented on the screen for 1.25 s and participants had to indicate their preferred item by pressing a button. The central fixation dimmed briefly when a response was registered. Snack stimuli shown here are for illustration purposes only. Participants viewed real branded items during the experiments. (b) Behavioral performance (red circles) and modelling results (black crosses). Participants’ average (N=21) reaction time (RT) and accuracy (top and bottom respectively) improved as the value difference (VD) between the alternatives increased. A sequential sampling model that assumes a noisy moment-by-moment accumulation of the VD signal fit the behavioral data well. (c) Average (N=21) model predicted evidence accumulation (EA) (black) and EEG activity (red) in the time window leading up to the response (on average, 600– 100 ms prior to the response), arising from a centroparietal electrode cluster (darker circles in the inset) that exhibited significant correlation between the two signals. Shaded error bars represent standard error across participants. (Reproduced with permission from (Pisauro et al. 2017))

Initial analysis of this data by (Pisauro et al. 2017) using the electrode whose raw EEG signal correlated best with the evidence accumulation model prediction, revealed a significantly positive cluster in posterior-medial prefrontal cortex (pMFC). In the original generalized linear regression model (GLM), raw EEG was used as the signal of interest along with three nuisance factors including visual stimulus onset, the value difference (of food items) and the (subject’s) reaction time (Figure S1).

As suggested previously (Wan et al. 2006) and based on physiological relationship between BOLD and energy consumption in a given region, we hypothesized that the instantaneous energy of a desired EEG electrode should be a more natural regressor against BOLD.

### Modelling the relation between EEG and BOLD

EEG and other extra-cellular measurements are the electric potentials associated with volume conduction of current dipoles arising from a bulk of activated neurons (Buzsáki et al. 2012). These dipoles arise from ion displacements across the cell membrane. The dissipated energy through these displacements would be proportional to the square of the membrane voltage, *V*^2^. Moreover, the electrical work performed by the active pumps can also be shown to be proportional to *V*^2^, since a linear relation between the pump’s current and the membrane voltage in a wide range of pump’s activity is previously reported (Nakao and Gadsby 1989). Therefore, the energy used in a voxel may be considered proportional to the square of “the electric potential or the magnitude of the current dipole” associated with this voxel. In particular, the electric potential for EEG electrode *i, e_i_* (*t*) can be written as the weighted sum of the dipole magnitudes from voxels across the brain;

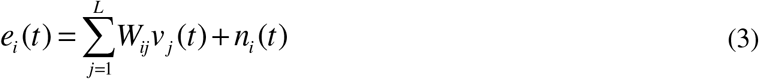

Where, L is the number of voxels (dipoles), *v_j_* (*t*) is the dipole magnitude associated with voxel *j*, *W_ij_* are the lead-field weights from voxels to the EEG electrodes and *n_i_* (*t*) is the noise present in EEG electrode *i*. The energy of this signal over a time interval of *T* equals:

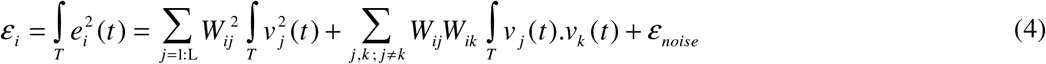

While the first summation in the right-hand side (R.H.S) of (4) is a weighted sum of the energies consumed in voxels, the second term is a summation of the voxels’ correlations. While the first term is strictly positive the second term can be suppressed due to positive and negative correlation across voxels. The noise term *n_i_* (*t*) is assumed to be orthogonal to the dipole time series. Thus, as a first order approximation, EEG energy can be considered as a weighted sum of dipole energy in each voxel which is the first term in eq (4). The error in this approximation can be quantified as the ratio of the second summation in eq (4) with respect to the whole sum as parameter z:

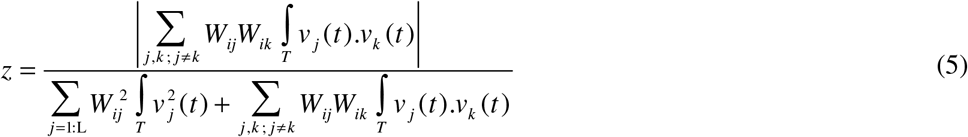

In order to verify the validity of this approximation, we ran simulations with various coherence levels between the dipole sources (see supplementary material). Simulations show that while this error term is an increasing function of correlation among dipole sources, nevertheless its maximum among the electrodes is less than 25% by average (Figure 2-a) even for coherences of up to 0.2 among voxels. The coherences below 0.2 are reported previously (Lehmann et al. 2012; Nentwich et al. 2020) and seem relevant even for patients with epilepsy and schizophrenia with high levels of synchrony among regions (Bowyer 2016).While the maximum of z among EEG electrodes provides the upper-bound of error, the expected value of error tends to be much smaller (less than 8%) in the same dynamic range of coherences between dipole sources (Figure 2-b). Figure 2-c also indicates that even for the highest correlation level, the maximum of z among electrodes lies most frequently between 10-20%.

**Fig. 2:**
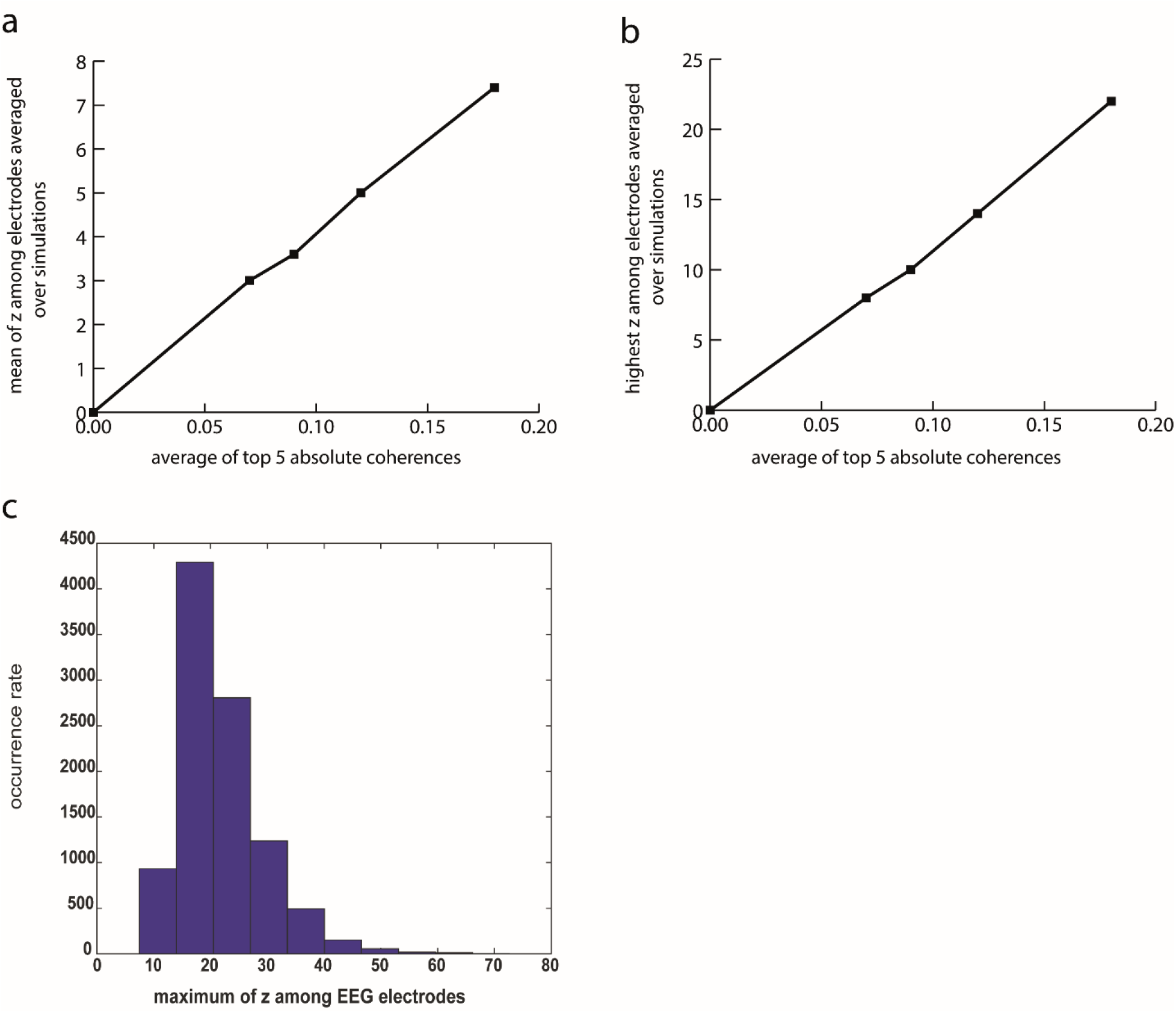
Effect of coherence between neuronal sources on the error percentage (‘z’) in approximating EEG energy as a sum of voxel BOLD values. A Gaussian random matrix sets the correlation between sources in each simulation trial. The coherence level of the sources is controlled by the variance of the Gaussian distribution and is measured by the average of the 5 pair of sources with the highest correlation. 10000 simulation trials were conducted for each coherence level. a) Effect of coherence level on the average (averaged across 10000 simulations) of maximum of ‘z’ among 63 EEG channels. b) Effect of coherence level on the mean of ‘z’ among 63 EEG channels averaged across simulations c) The histogram of “maximum of ‘z’ among EEG channels” for the highest coherence level simulated.

Therefore, we can approximately write:

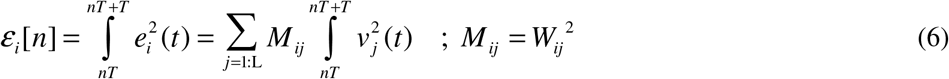

Obviously the route from the energy consumption in a voxel and the observed BOLD signal has to go through a couple of other steps including neurovascular coupling and blood vessel dynamics which subject this relationship to further nonlinearities and smoothing and can be modelled by detailed biophysical processes such as the Balloon model (Buxton et al. 1998). However, for simplicity here we only considered the simple hemodynamic function commonly used in the analysis of BOLD with GLMs. In this case convolving the two sides of (6) with the hemodynamic function we will have:

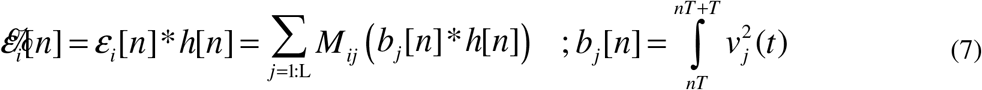

Since *b_j_* [*n*] * *h*[*n*] is assumed to be proportional to the BOLD signal of voxel *j* in the time volume *n*, the R.H.S of (7) is actually a weighted sum of the BOLD signals.

### EEG energy shows evidence accumulation in cortical regions involved in processing of food palatability

To examine whether “EEG energy” explains BOLD fluctuations over and above the 3 nuisance factors and the raw EEG, we used a step-wise GLM paradigm by first repeating the main GLM analyses in (Pisauro et al. 2017) (Table 1: GLM 1) and then regressing its residuals over the EEG energy regressor (Table 1: GLM 2). Interestingly, the activation map for the EEG energy (Figure 3; p-value < 0.05, cluster-corrected) was highly different from the activation map for the EEG (figure S2a) and showed significant positive correlation in the bilateral operculum, insula and the inferior somatosensory cortex as well as significant negative correlations across frontal, temporal, occipital and temporoparietal regions.

**Table 1.** inputs /outputs of the indexed GLMs

| GLM index | Signal to regress | Regressors |
| --- | --- | --- |
| GLM1 | BOLD | vstim – vd – rt – EEG |
| GLM2 | Residual of GLM1 | EEG energy |
| GLM3 | BOLD | vstim – vd – rt – vstim off – EEG |
| GLM4 | Residual of GLM3 | EEG energy |
| GLM5 | Residual of GLM3 | EEG pow3 |
| GLM6 | Residual of GLM4 | EEG pow 4 |
| GLM7 | BOLD | vstim (onset) – vd – rt – vstim boxcar |
| GLM8 | Residual of GLM7 | EEG |
| GLM9 | Residual of GLM7 | EEG energy |
| GLM10 | BOLD | vstim – vd – rt – vstim off |
| GLM11 | Residual of GLM10 | EEG (normalized) |
| GLM12 | Residual of GLM10 | EEG energy (normalized) |

**Fig. 3:**
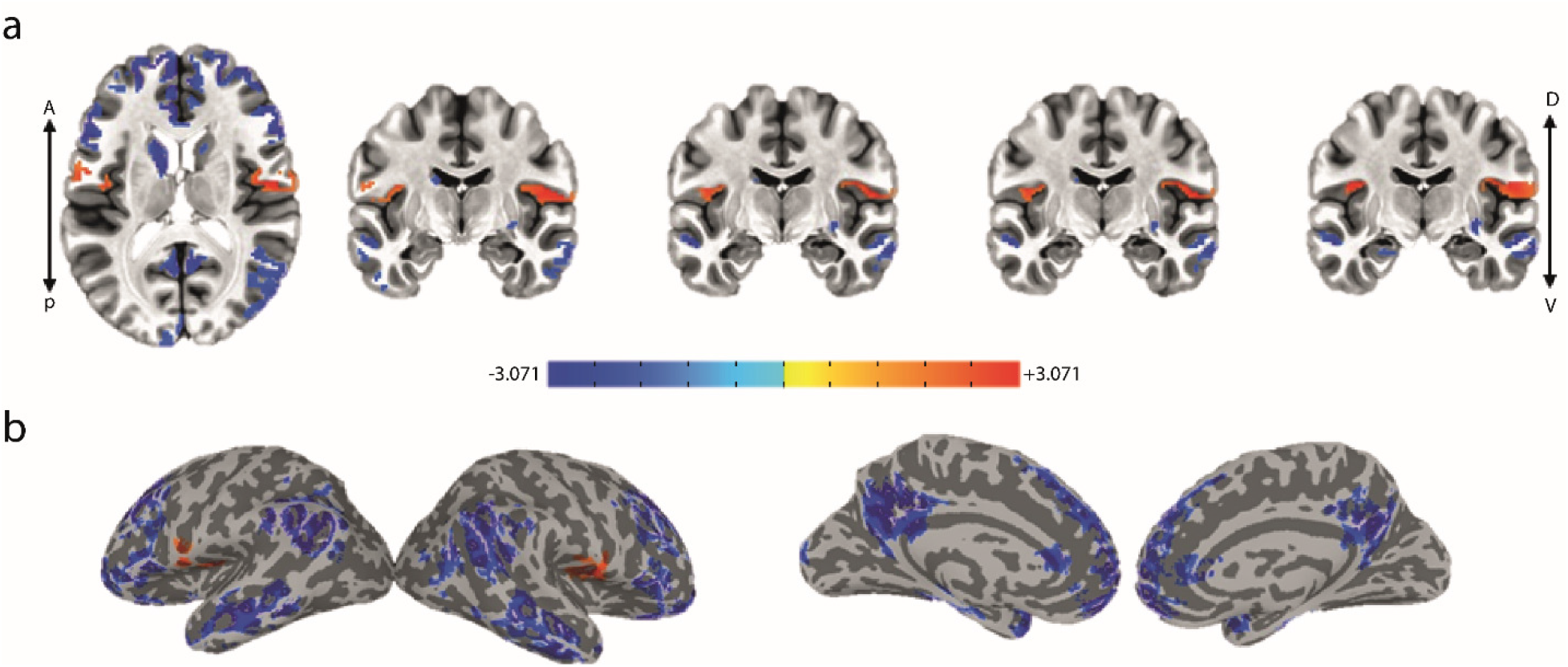
Group-average activation map (t-stats) for the “EEG energy” regressor in GLM2 showing activity in the bilateral insular, opercular and inferior somatosensory cortices (P-value < 0.05, cluster-corrected (right cluster=293, left cluster= 218 > threshold=136), a: axial and multiple coronal views, b: lateral and medial views on the inflated cortex.

Since in this task the process of evidence accumulation was fully overlapping with stimulus presentation, and in particular the fact that decision termination could be concurrent with stimulus offset, we repeated the previous analysis by adding ‘vstim-off’ regressor (Figure S1a) as an additional nuisance factor to ensure EEG related activations (especially those related to EEG energy) are not simply explained by temporal dynamics of sensory information on the screen. Indeed, it is shown that stimulus offset can evoke additional responses in the brain (Herdener et al. 2009; Mullinger et al. 2013, 2017). In this case, the first step GLM consisted of the four nuisance regressors plus EEG (Table 1: GLM 3). Then we regressed its residuals over the EEG energy (Table 1: GLM 4). Notably in this condition, the activation map for the EEG energy was similar to the case without inclusion of ‘vstim off’ (GLM2, Figure 3) but with positive correlation passing cluster correction threshold only in the right operculum and insula (Figure 4; p-value < 0.05, cluster-corrected, Table S1). This suggests that at least part of the positive activity seen in relation to EEG energy was not explainable by the ‘vstim-off’ regressor. On the other hand, inclusion of ‘vstim-off’, removed the previously reported activation in pMFC in relation to EEG altogether (Figure S2b) suggesting that the observed activity in pMFC to be due to visual offset rather than choice process per se.

**Fig. 4:**
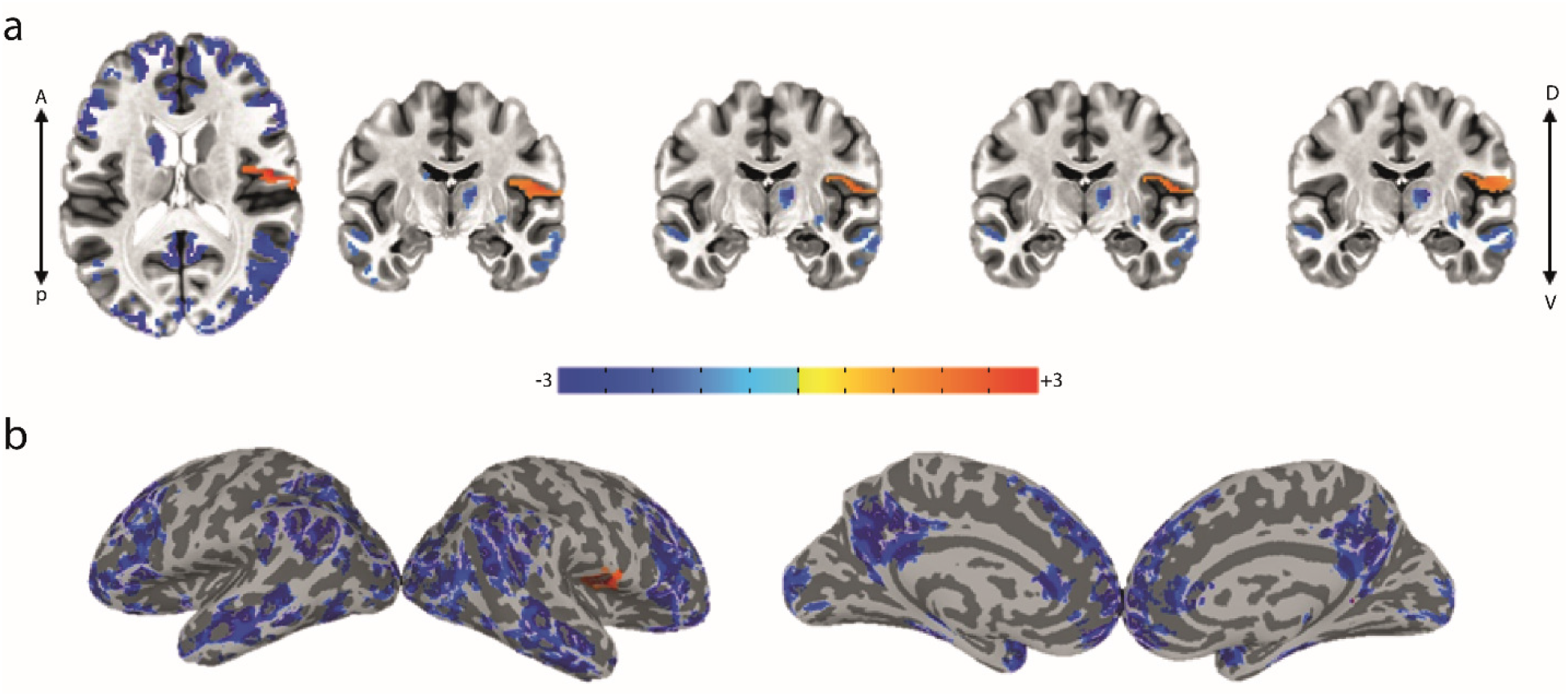
Group-average activation map (t-stats) for the “EEG energy” regressor in GLM4 showing activity in the right insular, opercular and inferior somatosensory cortices (P-value < 0.05, cluster-corrected (cluster=214 > threshold=121), a: axial and multiple coronal views, b: lateral and medial views on the inflated cortex.

**Fig. 5:**
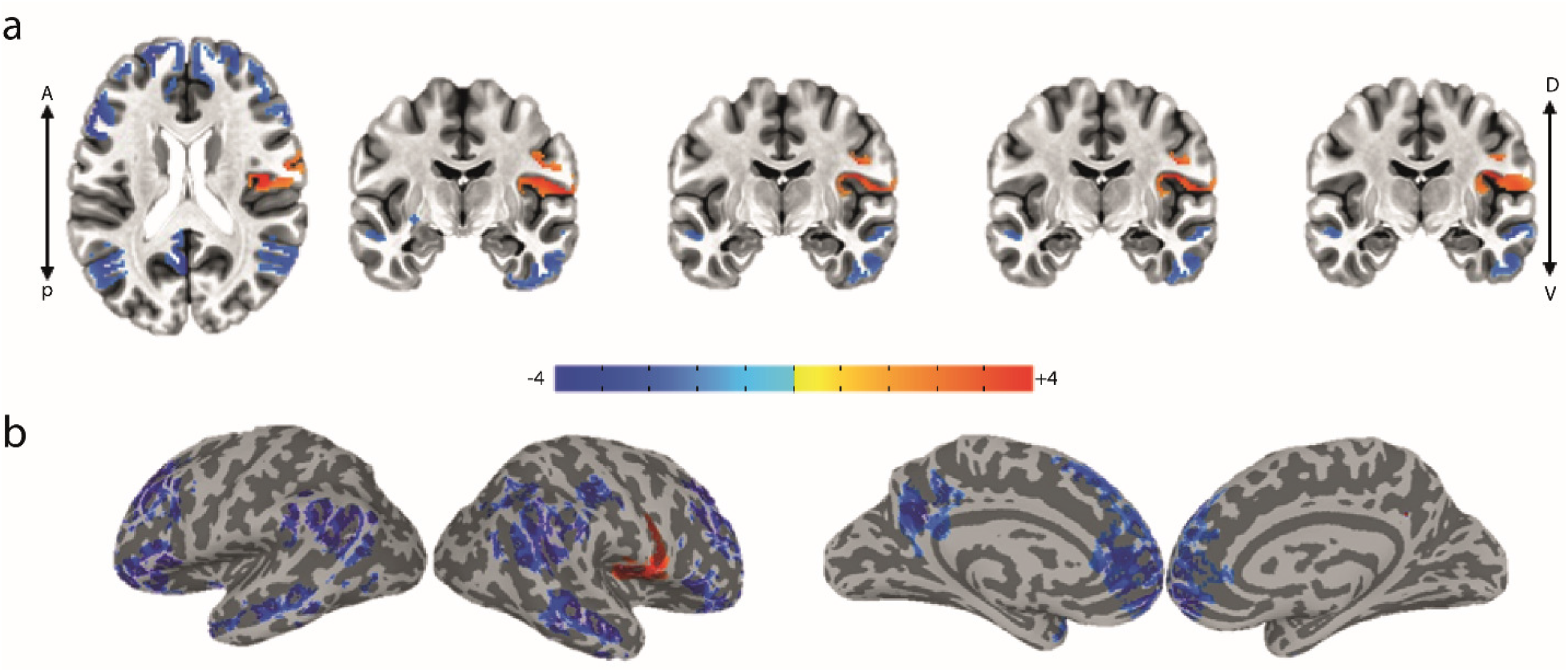
Group-average activation map (t-stats) for the normalized “EEG energy” regressor in GLM9 showing activity in the insular, opercular and inferior somatosensory cortices (P-value < 0.05, cluster-corrected (cluster=567 > threshold=295), a: axial and multiple coronal views, b: lateral and medial views on the inflated cortex.

Examination of activation maps for all four nuisance factors show a significant cluster for visual offset that overlaps with pMFC (Figure S3b). A qualitatively similar result was obtained if one were to use a boxcar for the duration of stimulus presentation as the nuisance factor instead of ‘vstim-off’ (GLM7). In this case again a significantly positive correlate in the right operculum and insula was observed for EEG energy (GLM8, Figure S4a). Once again, no significant cluster of activity related to EEG is observed in this case when stimulus duration is taken into account (GLM9, Figure S4b). These results suggest that the observed positive activations in insula is not explainable by stimulus dynamics and is most likely reflecting the process of value-based decision making.

The EEG power for different subjects may vary substantially due to multiple reasons such as intrinsic difference in neuronal activity level or due to differences in the lead-field gains. Therefore, it is often recommended to normalize the EEG signal of subjects in order to achieve a more reliable group-level inference (Cohen 2014). Here also we see a relatively wide dynamic range in mean EEG power across subjects (Figure S5). In order to make sure that such variability does not affect our main conclusions about significant positive activations observed in the right operculum, we performed another analysis with normalized EEG signals (Table 1: GLMS 11-12, see methods for details) on the residuals of regression model which included the four nuisance terms (Table 1: GLM 10). Results showed that EEG normalization reproduced the significant positive activation in the right operculum and insula in relation to EEG energy (GLM12, Figure 5; p-value < 0.05, cluster-corrected). Once again, we did not find any significant activity for the normalized raw EEG regressor in this case (GLM 11).

### Absence of significant cluster correlates with higher powers of EEG signal

Given the highly nonlinear mapping between electrical activity in a voxel and its BOLD signal such as those predicted by the Balloon model (Buxton et al. 1998), it is plausible that still higher order nonlinearities have to be considered in EEG-informed fMRI analysis. In order to check for possible higher order nonlinearities, we created regressors for higher powers of the EEG (power 3 and 4). To account for the multicollinearity among various powers of EEG, analysis of the power 3 of EEG was performed on the residual of first step GLM which included nuisance factors and the EEG regressor (GLM5). For power 4 of EEG, we did the regression on the residuals of EEG energy due to high correlation between power 2 (EEG energy) and power 4 (GLM6). Notably, no significant positive cluster of correlations with higher powers of EEG (powers 3 and 4) was found (Figure 6; p-value > 0.05, cluster-corrected). Some clusters of negative activation in power 3 of EEG was found close to prefrontal and temporoparietal areas (Table S1).

**Fig. 6:**
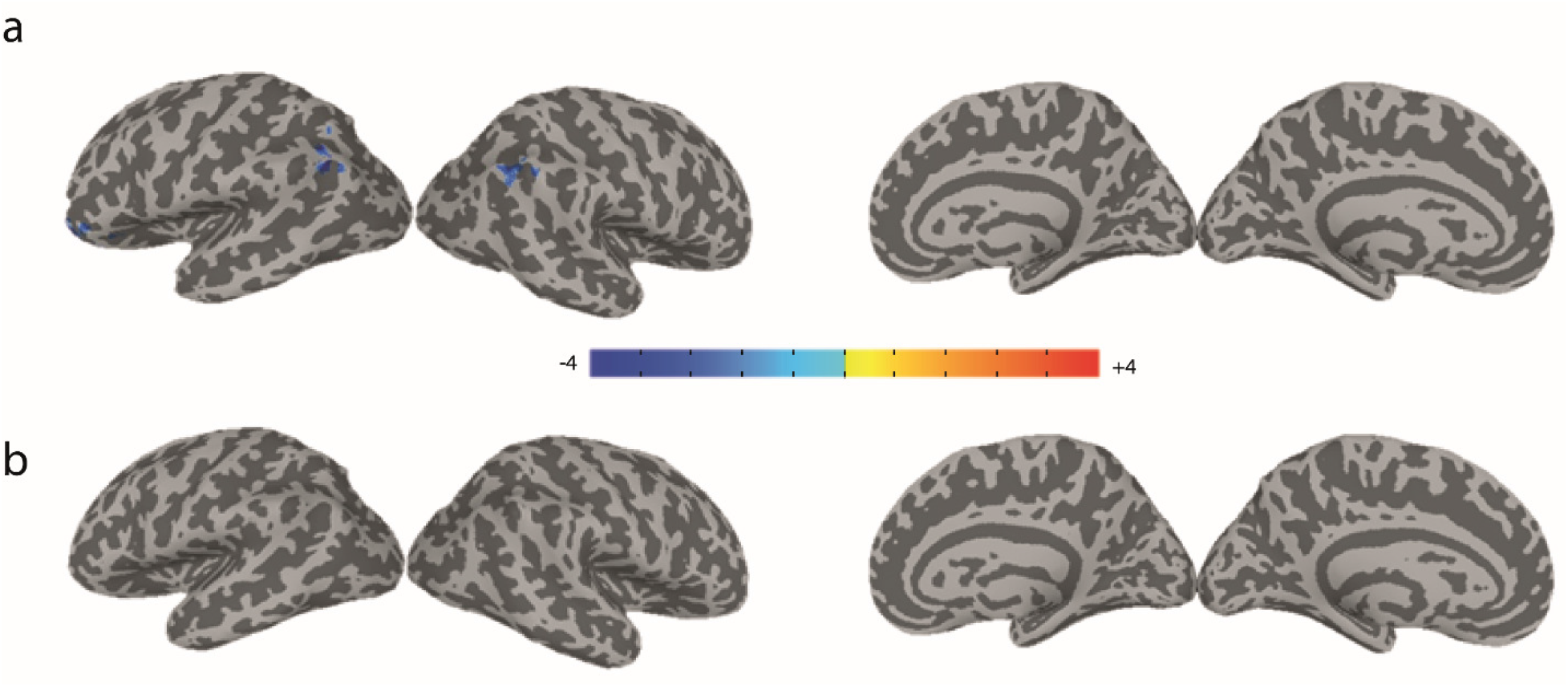
Group-average activation map (t-stats) for higher powers of EEG in GLM5 & GLM6; a) negative correlations with EEG pow3 (P-value < 0.05, cluster-corrected (clusters > threshold=121)), b: no significant correlations with EEG pow4 (P-value < 0.05, cluster-corrected (cluster threshold=71))

Finally, we did not observe any subcortical activations with any of the EEG-driven regressors used (raw EEG or any of its powers). The only notable subcortical activation was found in the amygdala in correlation with the “value difference” regressor (Figure S3-c).

## Discussion

EEG-informed fMRI analysis is a promising method in localizing fast cognitive processes in the brain such as the formation of a decision. Here, using simultaneous EEG-fMRI in a value-based decision-making task, revealed significant correlates of evidence accumulation in the insular and opercular cortices. This activity was uncovered by using “EEG energy” as the EEG-driven regressor and was missed if one were to use the raw EEG correlate of DV as was done previously (Pisauro et al. 2017). Here, we proved the relevance of EEG energy for BOLD theoretically, in agreement with the previous *experimental* evidence (Wan et al. 2006). Notably, despite the highly nonlinear nature of neurovascular coupling, we did not find any significant correlations with the higher powers of EEG (powers 3 and 4) in this task.

Given the multicollinearity of the experimental procedure at hand in which the decision formation and stimulus presentations were concurrent, one needed to make sure that the EEG related brain correlates were not due to stimulus onset, duration and offset. This problem was addressed by using step-wise GLMs which lets the nuisance regressors to describe as much as the variance in the BOLD as they can and leave the orthogonal components to be described by the regressor of interest in the second-step GLM (GLMs 2,4-6,8-9,11-12). Using this orthogonalization procedure, the “EEG energy” as the BOLD regressor revealed activity in the right operculum, insula and the inferior somatosensory cortex (Figures 3-5). Multiple parts of insula and operculum, including the anterior, middle and posterior insula as well as the frontal and parietal operculum extending to the inferior somatosensory cortex (area 3b) are reported to participate in taste and gustatory representations (Small 2010; Veldhuizen et al. 2011; Mai and Paxinos 2012) and form the gustatory cortex (GC). Some meta-analysis studies (Small 2010; Yeung et al. 2018) have distinguished the involvement of these diverse cites, in various aspects of taste and gustatory processing. The middle insula is reported for its role in coding the “pleasantness” aspect of taste and in attention to taste (Small 2010) as well as its participance in coding the affective value and quality of food regardless of its intensity (Yeung et al. 2018). The activity observed in the middle insula in our study, agrees well with the hypothesized role of insula in this task which is to evaluate the food’s pleasantness. It is conceivable for the activity in the GC to induce a gustatory imagery of the food items which are used in the evidence accumulation process for food choice. Interestingly, the insular and opercular regions have been also previously reported to be implicated in “gustatory *imagery*” (Kobayashi et al. 2004, 2011). Furthermore, the structural (Ogawa 1994; Ghaziri et al. 2018) and functional (Roy et al. 2009) connectivity between insula and amygdala and the fact that amygdala showed value-based activation in this task is in agreement with a possible value retrieval from amygdala.

Experimental evidence for a quadratic relation between the vascular input and the neural electrical sources estimated from EEG was provided previously (Wan et al. 2006) and used for studies involving epileptic patients (Murta et al. 2015; Abreu et al. 2018). Here, we extended the use of EEG energy to cognitive studies and argue from a theoretical stand-point that “EEG *energy*” should be a better correlate of BOLD response compared to EEG signal itself for the use in EEG-informed fMRI analyses. Consistent with this suggestion, correlations between the BOLD response and the power of EEG in the *alpha* band were reported in some studies especially in the resting state experiments (de Munck et al. 2009; Sato et al. 2010). Correlations between the BOLD response and various frequency bands of EEG in task-based experiments are also investigated (Scheeringa et al. 2009; Sato et al. 2010). A negative correlation between the *theta* power and BOLD response in the areas of the “default mode network, DMN” is reported in (Scheeringa et al. 2009). Actually, the negatively correlated regions with the EEG energy regressor in our study also highly overlap with the DMN network (Table S1), and this is plausible since the lower frequency bands of EEG (including *theta* band) dominate in EEG spectrum. These observed negative correlations with EEG energy during the decision making may suggest shutting down of these areas during value-based decision making.

The nonlinear nature of the neurovascular coupling could engender BOLD correlation with still higher powers of EEG signal. However, examining powers 3 and 4 of EEG signal in this study did not reveal any activations across the brain, suggesting “EEG *energy*” as a suitable and sufficient correlate of the BOLD response at least for this data.

Furthermore, we shall note that in spite of the observed significant activity in amygdala in correlation with the “value difference” regressor, there was no significant subcortical clusters in correlation with the EEG-driven regressors. This may indicate that the subcortical regions do not play a role in the process of evidence accumulation but only provide the needed inputs (value memory) for cortical regions responsible for decision making. On the other hand, this negative result may also be due the substantially lower signal-to-noise-ratio of the subcortical potentials on the EEG signal recorded on the scalp.

In summary, we conclude that due to the nonlinear relation between EEG and fMRI, “EEG energy” (or total power) proves critical for EEG-informed fMRI analysis. In particular, using EEG energy regressor in GLM of value-based decision-making revealed evidence accumulation activity in the operculum and insula. Activity in these regions as parts of the gustatory cortex indicates that gustatory imagery is likely to be used during the decision-making process for food choices and implicates cortical areas traditionally involved in palatability processing, in value-based decision making. Further investigations using electrophysiological techniques in human or non-human primates can help elucidate the exact dynamics of evidence accumulation in the gustatory areas during food choice.

## Acknowledgements

We would like to thank M. Andrea Pisauro and Elsa Fouragnan for comments and discussions on earlier versions of this manuscript.

## Code availability

All the codes related to this study can be accessed through a direct email to the corresponding author.

## Declarations

In this study, we used the free-access data from dataset and one can refer to (Pisauro et al. 2017) for more information about the ethics approval.

