## supplemental material for "Gustatory cortex is involved in evidence accumulation during food choice"

### **Supplementary material**:

#### Quantifying the error arising from ignoring voxel correlations in eq (4)

In order to quantify the amount of error arising from ignoring the second term in eq (4), we conducted multiple simulations for various correlation magnitudes between the sources using a real lead-field matrix. Specifically, we considered 20 active sources randomly distributed in the brain and produced the associated coherence matrix between these sources using a truncated Gaussian distribution. The overall correlation level between sources was controlled with the variance of this Gaussian distribution. We considered 4 levels of overall coherences and performed 10000 simulations for each level.

In each simulation, we recorded the average of the 5 highest correlation magnitudes between the sources (as the overall coherence level) as well as the maximum and average of z among the 63 EEG electrodes. Figure 2 shows the results of simulations which was explained in *results*.

#### Supplementary figures and table:

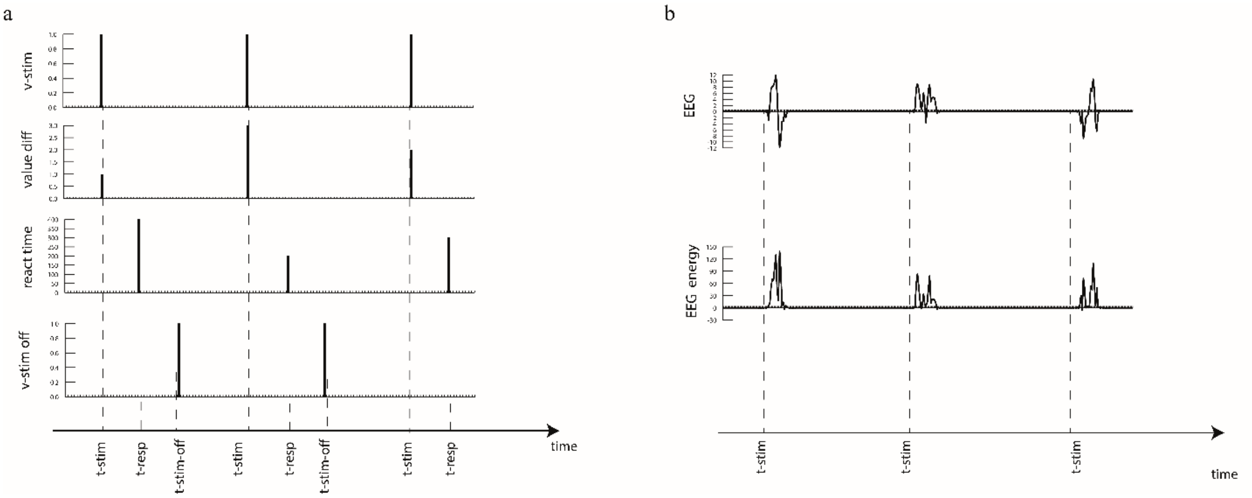


Fig. S1: The regressors used in fMRI analyses. a) The four nuisance regressors; the visual onset, the value difference, the reaction time and the visual offset regressors. b) The regressors of interest; the raw EEG and the EEG energy regressors

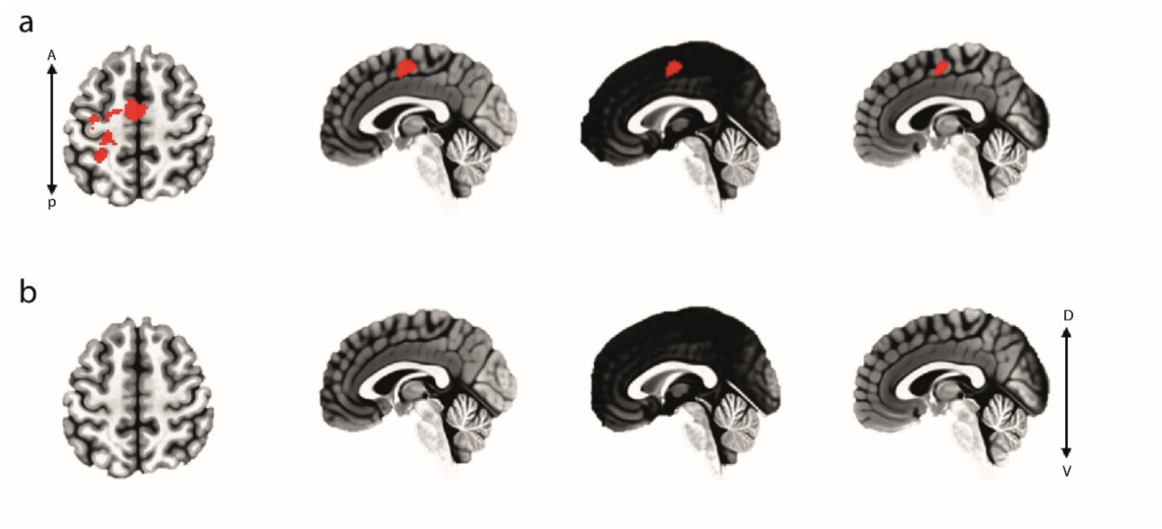


Fig. S2: Group-average activation map for the raw EEG regressor a) without considering “vstim-off” nuisance regressor as in GLM1 showing the activity in pMFC and premotor cortex (P-value < 0.05, cluster=1281> threshold=911) and in b) with considering “vstim-off” nuisance regressor, GLM3 (no significant activity with P-value < 0.05

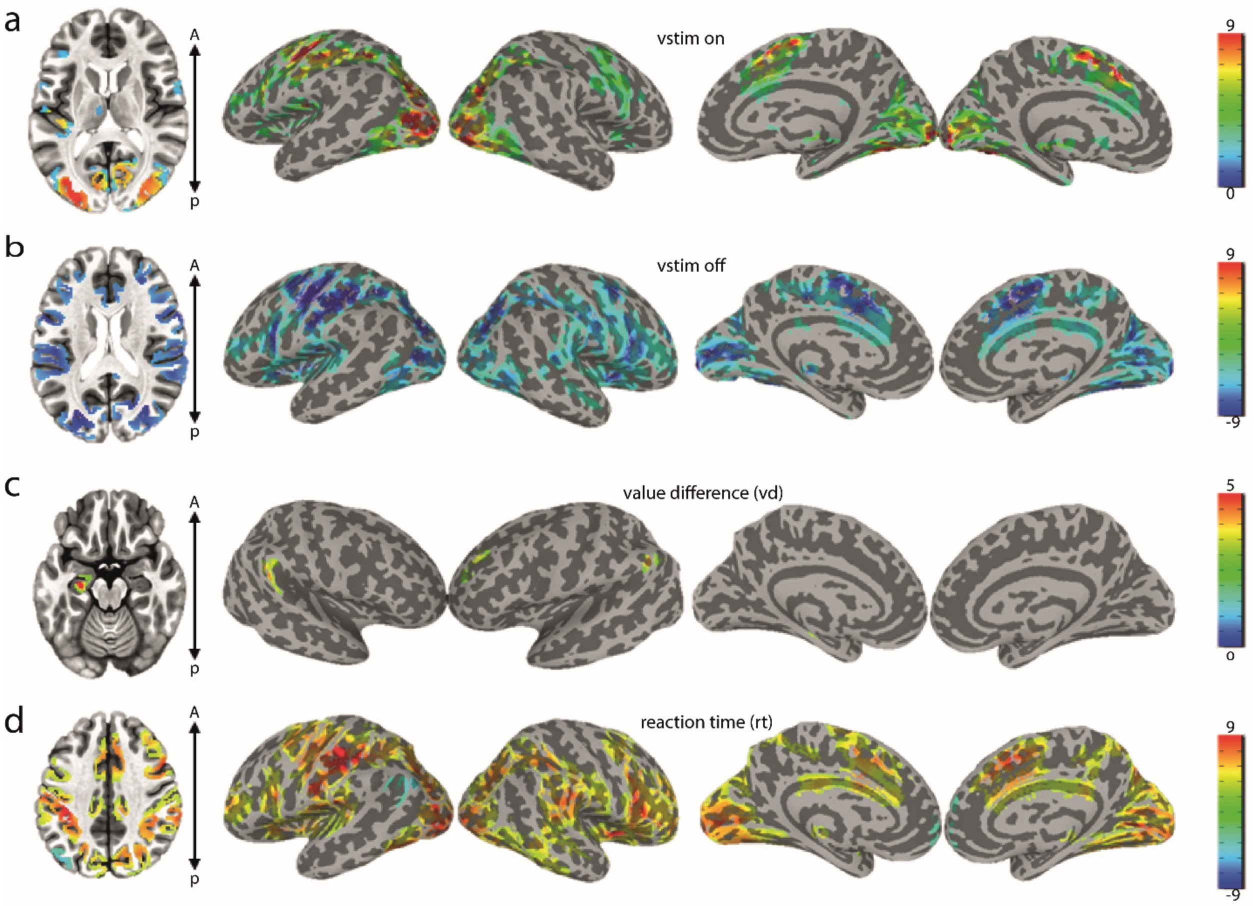


Fig. S3: Group-average activation map (t-stats) for the nuisance regressors all with P-value < 0.01 and cluster-corrected: a) visual onset regressor, b) visual offset regressor, c) value difference regressor and d) reaction time regressor

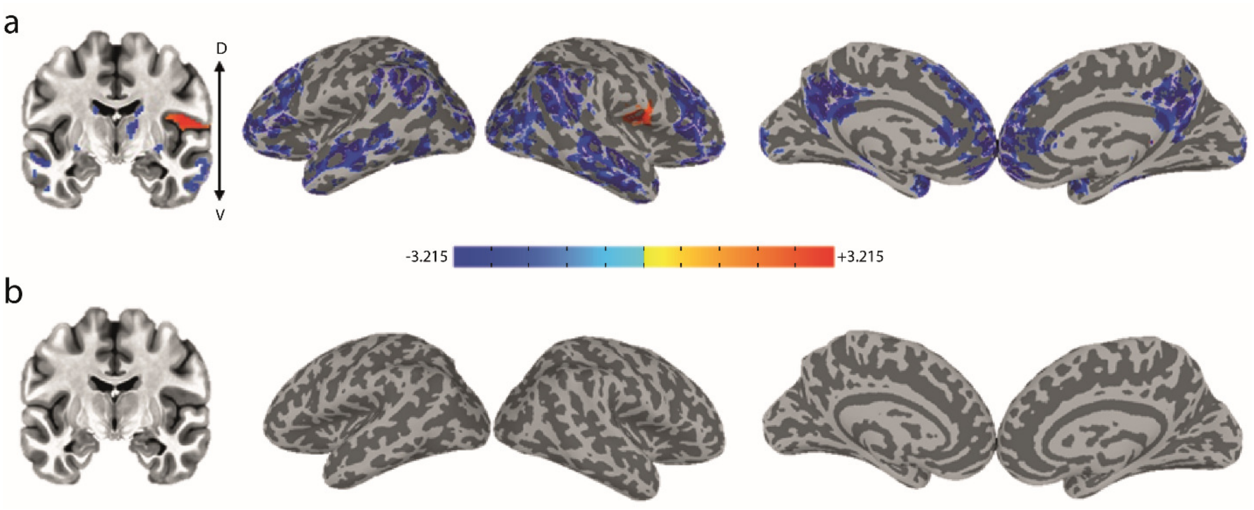


Fig. S4: Group-average activation map (t-stats) for a) the “EEG energy” regressor in GLM12 (P-value < 0.05, cluster-corrected, cluster = 219>threshold=138) and b) the raw EEG regressor in GLM11 with no significant (P-value < 0.05) activity.

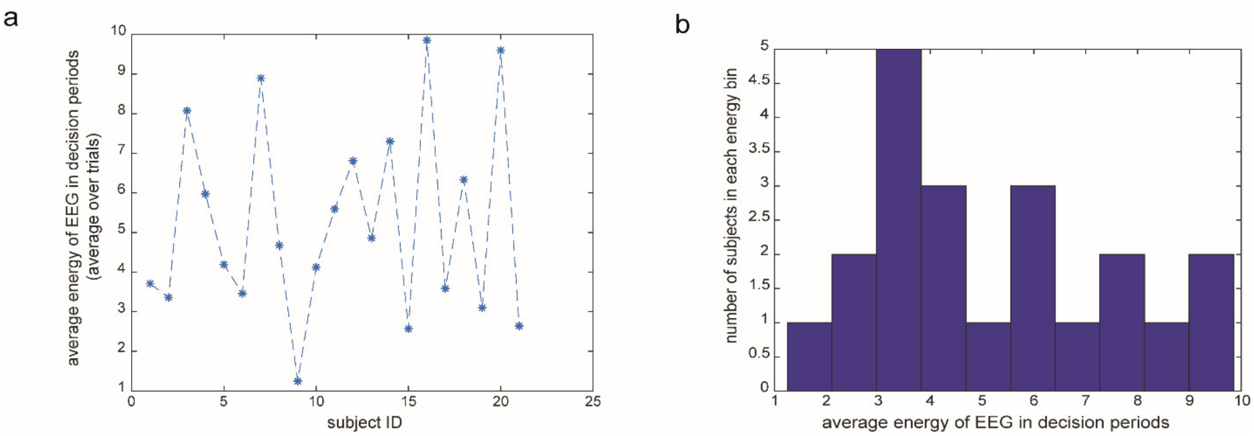


Fig. S5: Average energy of the best electrode EEG in decision periods for different subjects showing a large dynamic range. a) Average of “EEG energy” for each subject b) Histogram of average “EEG energy” across all subjects.

Table S1, table of activated regions with the EEG-driven regressors

| region/cluster | #(voxels) | hemisphere | peak X | peak Y | peak Z |
|---|---|---|---|---|---|
| Energy(+) | | | | | |
| | 214 | right | -42 | 10.0 | 22.0 |
| Energy(-) | | | | | |
| | 10945 | right | -18 | -56 | 30 |
| | 3351 | right | -48 | 54 | 56 |
| | 2157 | Left | 58 | 54 | 36 |
| | 2048 | Left | 30 | -14 | -34 |
| | 1668 | right | -52 | 22 | -12 |
| | 928 | Left | 12 | 44 | 30 |
| | 872 | Left | 28 | 90 | 22 |
| | 791 | right | -4 | 44 | 34 |
| | 705 | Left | 38 | 64 | -2 |
| | 683 | right | -34 | 68 | -26 |
| | 603 | right | -22 | -12 | 0 |
| | 528 | right | -44 | -10 | -20 |
| | 488 | Left | 28 | -10 | -18 |
| | 257 | Left | 8 | -4 | 6 |
| | 150 | right | -14 | 12 | 12 |
| | 148 | right | -30 | 46 | -32 |
| EEG pow3 (-) | | | | | |
| | 275 | Left | 40 | -50 | -10 |
| | 140 | right | -46 | 68 | 48 |
| | 135 | Left | 40 | 66 | 48 |
